# Differentially expressed tRNA-derived fragments in bovine fetuses with assisted reproduction induced congenital overgrowth syndrome

**DOI:** 10.1101/2022.09.28.509974

**Authors:** Anna K. Goldkamp, Yahan Li, Rocio M. Rivera, Darren E. Hagen

## Abstract

**Background:** As couples struggle with infertility and livestock producers wish to rapidly improve genetic merit in their herd, assisted reproductive technologies (ART) have become increasingly popular in human medicine as well as the livestock industry. Utilizing ART can cause an increased risk of congenital overgrowth syndromes, such as Large Offspring Syndrome (LOS) in ruminants. A dysregulation of transcripts has been observed in bovine fetuses with LOS, which is suggested to be a cause of the phenotype. Our recent study identified variations in tRNA expression in LOS individuals, leading us to hypothesize that variations in tRNA expression can influence the availability of their processed regulatory products, tRNA-derived fragments (tRFs). Due to their resemblance in size to microRNAs, studies suggest that tRFs target mRNA transcripts and regulate gene expression. Thus, we have sequenced small RNA isolated from skeletal muscle and liver of day 105 bovine fetuses to elucidate the mechanisms contributing to LOS. Moreover, we have utilized our previously generated tRNA sequencing data to analyze the contribution of tRNA availability to tRF abundance.

**Results:** 22,289 and 7,737 unique tRFs were predicted in the liver and muscle tissue respectively. The greatest number of reads originated from 5’ tRFs in muscle and 5’ halves in liver. In addition, mitochondrial (MT) and nuclear derived tRF expression was tissue-specific with most MT-tRFs and nuclear tRFs derived from Lys^UUU^ and iMet^CAU^ in muscle, and Asn^GUU^ and Gly^GCC^ in liver. Despite variation in tRF abundance within treatment groups, we identified differentially expressed (DE) tRFs across Control-AI, ART-Normal, and ART-LOS groups with the most DE tRFs between ART-Normal and ART-LOS groups. Many DE tRFs target transcripts enriched in pathways related to growth and development in the muscle and tumor development in the liver. Finally, we found positive correlation coefficients between tRNA availability and tRF expression in muscle (R=0.47) and liver (0.6).

**Conclusion:** Our results highlight the dysregulation of tRF expression and its regulatory roles in LOS. These tRFs were found to target both imprinted and non-imprinted genes in muscle as well as genes linked to tumor development in the liver. Furthermore, we found that tRNA transcription is a highly modulated event that plays a part in the biogenesis of tRFs. This study is the first to investigate the relationship between tRNA and tRF expression in combination with ART-induced LOS.

## Introduction

Assisted reproductive technologies (ART) are described as treatments that manipulate reproduction to increase chances of conception and encompasses a wide array of procedures such as *in vitro* fertilization (IVF), intracytoplasmic sperm injection (ICSI), and embryo transfer [1]. ART is often used to increase genetic gain and advance reproductive potential, and its use has rapidly increased in beef and dairy cattle populations [2]. In fact, a 2017 report showed a dramatic shift in worldwide embryo production, in which significantly higher numbers of bovine embryos are now produced *in vitro* compared to *in vivo* [3]. According to the International embryo technology society (IETS) newsletter, a record of more than 1.5 million *in vitro*-conceived bovine embryos were produced or collected in 2020 alone [4]. Although *in vitro* embryo production has quickly become the preferred technique globally, it is important to consider the effects of *in vitro* procedures on genomic output.

Several studies have investigated the link between ART use and the increased occurrence of congenital overgrowth syndromes, such as Beckwith-Wiedemann syndrome (BWS) in humans and Large Offspring Syndrome (LOS) in ruminants [5–9]. LOS is often characterized by overgrowth, tongue enlargement, and abdominal wall defects [5, 10, 11]. BWS shares clinical features with LOS and is also associated with an increased risk of liver tumors (hepatoblastoma) [12]. Livestock are often bred for economically beneficial characteristics related to production, making LOS an issue for breeders and a source of economic loss for producers. Due to their large size, LOS offspring have an increased chance of dystocia (difficult birth) which can result in death of the calf and/or dam [13]. In addition to cow and calf mortality, dystocia can result in financial losses associated with decreased milk production and fertility, and an increased likelihood of health issues (e.g. respiratory and digestive disorders, uterine disease, mastitis) [14–17]. However, the mechanism of ART-induced fetal overgrowth remains poorly understood.

Our previous work has detected dysregulation of transcripts and differentially methylated regions (DMRs) in LOS, and some of these regions resulted in dysregulation of imprinted loci (genes expressed in a parent-specific fashion) [18, 19]. We also have shown that DNA methylation is associated with a very small percent of gene misregulation in LOS individuals, suggesting other factors may be influencing gene regulation [20]. Therefore, there is still a lack of clarity in diagnosis due to the variation in molecular basis and presence of major clinical symptoms. Due to their crucial role in protein synthesis, our recent study investigated tRNA expression within skeletal muscle and liver in LOS. This study revealed differential expression of tRNA genes as well as tissue- and treatment-specific tRNA transcripts with unique sequence variations [21].These findings as well as the discovery of small non-coding RNAs derived from tRNAs, led us to consider the role of tRNA-derived fragments (tRFs) in LOS. To date, no study has examined the relationship between bovine tRNA expression and their processed regulatory products.

During tRNA maturation, the 5’ leader and 3’ trailer sequence of precursor tRNAs (pre-tRNAs) is cleaved by RNase Z and RNase P [22, 23] **(Figure 1)**. Following the addition of a 3’ CCA tail and enzymatic splicing, the mature tRNA is actively transported through the nuclear pore complex. Mature tRNAs may be cleaved through a Dicer-dependent or Dicer-independent pathway and several classes of tRFs are produced based on the tRNA cleavage position: 5’ tRFs, 3’ tRFs, internal tRFs (i-tRFs; internal fragments spanning anywhere within the tRNA), 5’ halves, and 3’ halves **(Figure 1)**. Generally, 5’ tRFs, 3’ tRFs, and i-tRFs are 16-26 nt, whereas 5’ and 3’ halves are 27-36 nt [23]. Initially considered to be random tRNA degradation products, growing evidence indicates that tRFs are an emerging class of non-coding RNAs with implications in multiple biological processes, namely regulation of protein translation [24, 25]. There are several suggested mechanisms of translational inhibition, such as disrupted ribosomal interactions through mRNA competition [26], displacement of initiation factors necessary for translation [27], and recruitment of RNase Z to cleave target mRNAs [28]. Other studies suggest that stress granules may be formed in response to tRF-mediated inhibition of protein synthesis, which can reduce apoptosis in cancer cells [29, 30]. The primary pathway that is frequently suggested is similar to microRNAs, in which the tRF is loaded into an RNA-induced silencing complex (RISC) to target partially complementary mRNA [31–33]. Various tRF types have been identified in plants, humans, and cattle with some acting as promoters of metastasis or aiding in homeostasis in humans [34, 35], and others for example have been reported to respond to nutritional deficiency in *Arabidopsis* [36] or Bovine Leukemia Virus in cattle [37]. Considering the alterations in tRNA expression and the dysregulation of mRNA transcripts in LOS, these tRNA genes may be selectively transcribed to give rise to unique tRF subtypes capable of targeting transcripts related to growth and/or liver tumor development. Furthermore, the diverse functions of tRFs across health states indicates a possible role in syndrome development.

**Figure 1.**
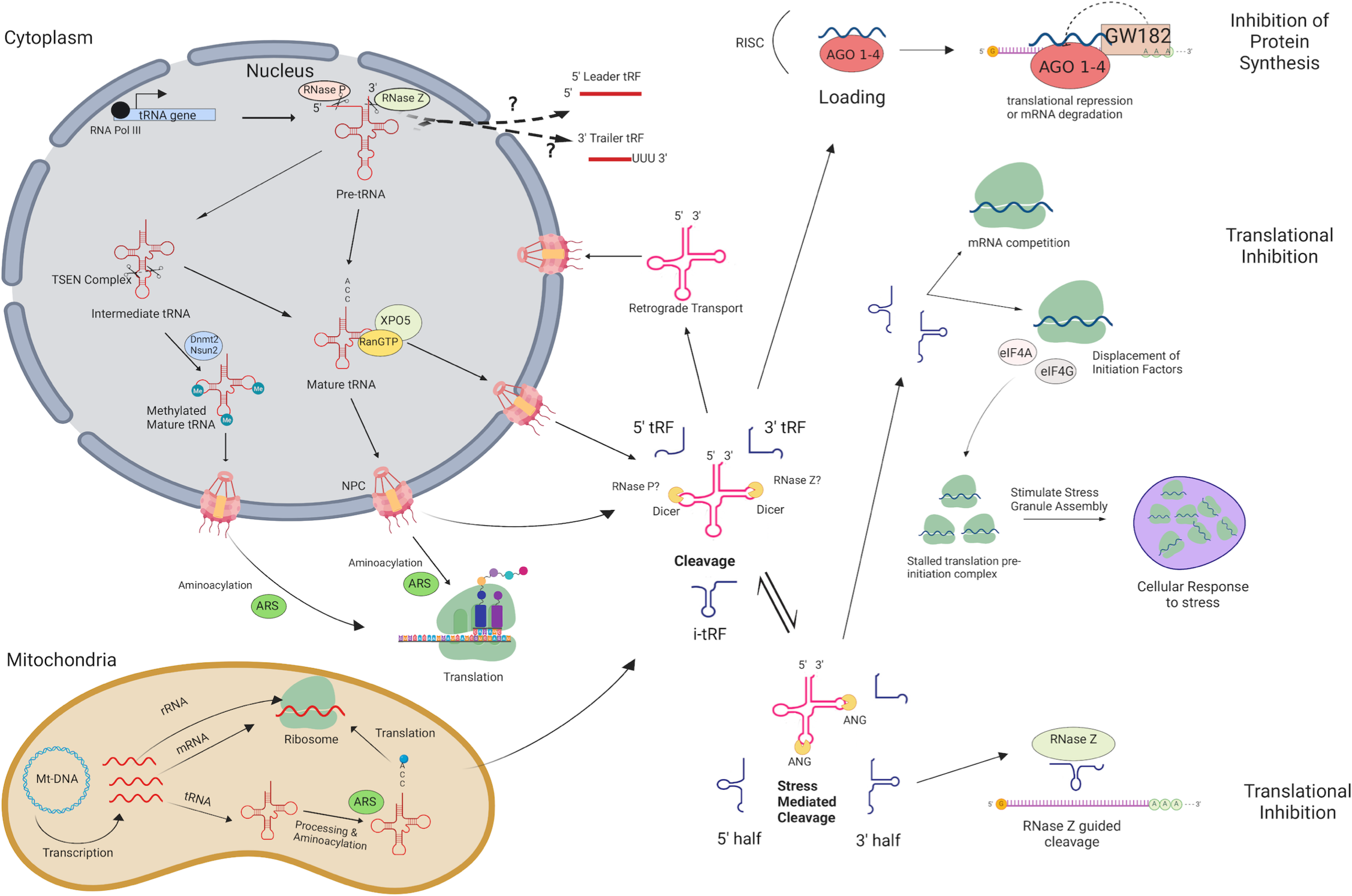
tRNA & tRF biogenesis pathway and silencing mechanisms in eukaryotes. tRNAs and tRNA-derived fragments (tRFs) are produced through this pathway in eukaryotes. Arrows indicate each step and help visualize how mRNA transcripts are targeted for translational repression or degradation, as well as how they influence stress response. There are differences in cellular ribonucleases used for cleavage in bacteria and yeast that are not shown. RNA POL III: RNA Polymerase III; TSEN Complex: tRNA Splicing Endonuclease Complex; AGO1-4: Argonaute 1-4; XPO5: Exportin-5; RanGTP: GTP-bound Ras-related nuclear protein; Nsun2: NOP2/Sun RNA methyltransferase family member 2; Dnmt2: DNA methyltransferase 2; NPC: Nuclear Pore Complex; ARS: aminoacyl-tRNA synthetase; ANG: Angiogenin; Mt-DNA: Mitochondrial DNA; RISC: RNA-induced silencing complex; eIF4A: eukaryotic translation initiation factor 4A; eIF4G: eukaryotic translation initiation factor 4G. Figure created with BioRender.com.

In this study, we performed small RNA sequencing on skeletal muscle and liver samples collected from day 105 artificial insemination-conceived fetuses (AI-Control), ART-conceived bovine fetuses with a body weight above the 97^th^ percentile relative to Control-AI (ART-LOS), and ART-conceived bovine fetuses with a body weight below the 97^th^ percentile (ART-Normal). In addition, previously generated tRNA sequencing data was used to compare the expression of mature tRNAs and their processed regulatory products [21]. We detected differentially expressed tRFs due to method of conception (AI vs ART) as well as syndrome development (ART-Normal vs ART-LOS). Our results indicate that tRNA expression is highly dynamic based on tissue type and syndrome development. This brings the possibility that some tRNA expression can act as a means of tRF production in order to regulate gene expression. This study contributes insights on the mechanisms of tRF biogenesis and their role in targeting transcripts related to growth and development.

## Materials and Methods

### Animals and RNA isolation

Day 105 *Bos taurus indicus* (B. t. indicus; Nelore breed) × *Bos taurus taurus* (B. t. taurus; Holstein breed) F1 fetal conceptuses were previously generated by us [38]. Tissues were flash frozen in liquid nitrogen and stored at –80° until RNA extraction. Total RNA was extracted from skeletal muscle and liver tissues of F1 hybrid controls (artificial insemination; Control-AI), *in vitro* produced ART-Normal (similar weight as controls), and *in vitro* produced ART-LOS (body weight greater than 97^th^ percentile relative to controls) using TRIzol Reagent (Invitrogen, Carlsbad, CA) following the manufacturer’s instructions. Quality and concentration of the RNA samples was assessed using the Agilent TapeStation RNA ScreenTape (Agilent, Santa Clara, CA) and RNA integrity numbers (RIN) for all samples were ≥ 7.4.

### Library preparation and sequencing

Small RNA library preparation was done using the TruSeq® Small RNA Library Preparation Kit (Illumina, Inc., San Diego, CA) and following the manufacturer’s instructions. 1 μg of total RNA and was briefly ligated to 3’ RNA adapters with ligation buffer, RNase Inhibitor, and T4 RNA Ligase 2. After the addition of stop solution, 5’ RNA adapters were also ligated with T4 RNA Ligase 2. Reverse transcription was performed with each adapter-ligated RNA library to produce cDNA constructs. Each resulting cDNA library was amplified via Polymerase Chain Reaction (PCR). A unique RPIX was used for each library sample for multiplexed sequencing and analysis. Following PCR and before cDNA construct purification, each library was run on a High Sensitivity DNA chip (Agilent, Santa Clara, CA) with expected peaks of approximately 140-160 bp. The pooled libraries were resolved on a 6% Novex TBE PAGE gel (polyacrylamide gel) and a size selection of 140 to 180 bp (predicted size of tRNA fragments and adapters) was performed on the gel. The purified and pooled libraries were sequenced using Illumina NextSeq 500 System High-Output Kit (Illumina, Inc., San Diego, CA) and conducted by the OSU Microarray Core Facility. All lanes were sequenced at the same time to prevent a batch effect. The liver small RNA-seq data was provided from our previous study (GEO database accession # GSE117015) [38] and was sequenced using Illumina NextSeq 500 System High-Output Kit, the same library preparation 158 kit and sequencing platform as the skeletal muscle tRFs, by the University of Missouri-Columbia DNA core facility.

### Processing and alignment of small RNAseq data

The raw sequence reads were filtered using the fastq-mcf command from ea-utils (version 0.148d4) in order to remove the TruSeq Small RNA adapter sequence (TGGAATTCTCGGGTGCCAAGG) [39]. The adapter trimmed reads were then quality trimmed using SolexaQA++ (version 3.1.6) dynamictrim utility with a Phred cut off score of 19 [40]. The quality trimmed reads were kept if they had a length of at least 13 bp or greater and were sorted using the SolexaQA++ lengthsort utility. The resulting reads were then mapped against the bovine genome ARS-UCD1.2 using the MINTmap pipeline in order to predict tRNA fragments from the small RNA-seq data [41, 42]. MINTmap aligned reads to a look up table that contains unique tRF sequences ranging from 16 to 50 bp that are exclusively located in regions associated with annotated tRNA genes. The reads that mapped to bovine tRFs were measured with the default setting of MINTmap, which allows no mismatches, no insertions, and no deletions and also analyzes the whole genome to retrieve all possible alignments [41]. Additionally, MINTmap outputs the parental tRNA source(s) that the tRF is potentially derived from, the tRF sequence, tRF subtype and the unique MINTplate associated with the tRF. Only the exclusive tRF expression output of unnormalized reads was used for data analysis.

### Differential Expression analysis

Non-linear full quantile normalization was used with the betweenLaneNormalization function on CPM transformed read counts using EDAseq v2.24.0 in order to produce PCA and RLE plots. Principal component analyses (PCA) and relative log expression (RLE) plots were created with the plotPCA and plotRLE function of the EDASeq package, respectively. Only tRFs that had at least 5 counts per million in all of the control, or all of the ART-normal, or at least 2 ART-LOS were considered moderately expressed and kept for DE analysis. EdgeR v3.24.3 was used to conduct a differential expression analysis and the trimmed mean of M values method (TMM) of EdgeR was used for normalization [43]. A likelihood ratio test was conducted using the glmLRT function of edgeR in order to identify differential expression in skeletal muscle and liver (Control-AI vs ART-normal, ART-normal vs ART-LOS, and Control-AI vs ART-LOS). Differentially expressed tRFs were defined as those with a false discovery rate (FDR) of ≤ 0.05. Heatmaps of differentially expressed tRFs were created for skeletal muscle and liver tissue to graphically represent gene expression. The normalized read counts were transformed into moderated log-counts per million and heatmaps were produced using RColorBrewer v1.1-2 and the heatmap.2 function of the gplots package v3.0.1.1.

### Target prediction

RNA-seq data for Control and ART-LOS individuals from our previous study was retrieved from NCBI Gene Expression Omnibus (GEO) (accession # GSE63509) [44]. RNA-seq data was used to predict potential gene candidates targeted by DE tRFs. Differential expression analysis of the RNA-seq data was done using the same method previously described for the tRF analysis.

All DE tRFs that were identified between Control vs ART-Normal, ART-Normal vs ART-LOS, and Control vs ART-LOS were analyzed for target prediction. The 3’ UTR sequences of all expressed protein-coding genes in the ARS-UCD1.2 bovine genome were obtained from Ensembl Release 98 [45]. miRanda v3.3a is a program commonly used for miRNA target prediction and was used for DE tRF target prediction in this study [46, 47]. The 3’ UTR sequences of the protein coding genes were used as a reference for alignment of the DE tRF sequences with a binding score cutoff of ≥ 150 and an energy cutoff of ≤ –20 [47]. Since it has been proposed that tRFs may target transcripts that are only partially complementary, unlike miRNAs, the strict parameter was not used and a partially complementary seed sequence was allowed [48, 49]. Predicted tRF targets were compared with the DE transcripts obtained from analysis of RNAseq mentioned previously. Downregulated mRNA targets were overlapped with targets of upregulated tRFs and vice versa for each treatment comparison.

### Functional Enrichment Analysis

Enrichment analysis was done using the list of candidate gene targets of DE tRFs for each treatment comparisons and all expressed protein coding genes used as the background gene set. Gene set enrichment analysis (GSEA) of the GO terms was performed using Fisher’s exact test as implemented in R package topGO v2.42.0 [50]. KEGG pathway enrichment analysis was performed using the Wilcoxon rank-sum tests via the R package KEGGREST v1.30.1 [51]. Human Phenotype Ontology (HPO) enrichment analysis was done using the g:Profiler web server with p-values corrected by the g:SCS threshold significance criterion. [52]. For the identified candidate target genes, we used mouse mutant phenotype information and performed a mammalian phenotype enrichment analysis with the Fisher’s Exact test implemented by MamPhEA [53]. Enrichment results with a p-value of ≤ 0.05 were classified as significant. Dot plots depicting enrichment results were created with ggplot2 package v3.2.1.

### YAMAT-seq data

Mature tRNA sequencing data from our previous study was used to compare tRNA and tRF levels [21]. MINTmap was used to provide all possible parental tRNA sources of each tRF [41, 42]. In order to evaluate the relationship between parent tRNA expression and tRF abundance, both YAMAT-seq and small RNAseq data were then merged. In an effort to not exclude any parent tRNA predicted by MINTmap, the counts for each tRF were divided by the number of parental tRNAs it was predicted to be derived from and were then log transformed. If there was no detected expression in both the parental tRNA and the tRF in any treatment group, the tRNA species was not included in the scatter plot. Scatter plots were made to show tRNA and tRF expression relative to the tRNA species with ggplot2 package v3.2.1 and by tissue type to calculate Pearson’s correlation coefficient with ggpubr package v0.2.4.

## Results and Discussion

### Small RNA sequencing

In order to understand tRF expression in bovine fetuses with congenital overgrowth syndromes, we performed small RNA sequencing to generate tRF expression profiles in skeletal muscle and liver. This resulted in an average of 10,750,864 (76.7%) and 9,074,956 (86%) reads retained per sample for muscle and liver respectively **(Table 1)**. Adapter and quality trimmed reads were aligned to the ARS-UCD1.2 bovine reference genome using the MINTmap pipeline in order to predict tRNA fragments from the small RNA-seq data. A total of 936,898 and 2,854,063 reads exclusively mapped to tRFs in the skeletal muscle and liver. A lower proportion of retained reads were mapped due to MINTmap’s strategy: only exact matches are allowed, one sequence is counted once no matter how often it appears within the genome, and only tRFs that map exclusively to genomic tRNA locations are counted. Therefore, we excluded ambiguous reads that mapped to locations both within and outside of tRNA loci in order to prevent false positives.

### Detection of tRNA-derived fragments

Five subtypes of mapped tRFs were predicted in muscle and liver datasets: 5’-tRF, 3’-tRF, i-tRF, 5’ half, and 3’ half. A total of 22,289 unique tRFs were predicted in the liver tissue and 7,737 unique tRFs were predicted in the muscle tissue **(Supplementary Table 1)**. The larger number of predicted tRFs in liver could be a result of high transcriptional activity in the liver tissue. Our recent tRNA study detected a greater number of tRNA genes expressed in liver compared to muscle (487 vs 474), which could contribute to changes in the tRF profile of each tissue [21]. The liver acts as a key player in nutrient metabolism and detoxification, which could result in excess transcripts in an effort to effectively regulate metabolic homeostasis. In fact, the human liver transcriptome has been described to have increased complexity and significant variability in transcript expression [54, 55]. We included a filtering step, in which tRFs with counts present in any two individuals within a tissue (n = 10) were classified as expressed and kept for analysis. This filtration step yielded a total of 13,231 tRFs in the liver and 3,508 tRFs in the muscle. Out of the 13,231 expressed tRFs in the liver, the distribution of tRFs by subtype are as follows: 11,102 i-tRFs, 1,492 5’-tRFs, 305 5’ halves, 294 3’-tRFs, and 38 3’ halves. Out of the 3,508 expressed tRFs in the muscle, there were 2,748 i-tRFs, 687 5’-tRFs, 48 5’ halves, and 25 3’-tRFs. **(Supplementary Table 1**). i-tRFs were the most common of the list of predicted tRFs in both tissues. i-tRFs arise from a variety of positions and may be derived upstream, within, or downstream of the anticodon loop [41]. This creates an opportunity for more than one i-tRF to be processed from a single tRNA molecule. Despite most of the predicted tRF species being of the i-tRF subtype in both tissues, the distribution of reads derived from a particular subtype was tissue-specific. The largest portion of reads were derived from 5’ tRFs in the muscle and 5’ halves in the liver **(Figure 2A)**. These results are consistent with previous studies in human and mouse which reported that hematopoietic tissues, such as the liver, have greater expression of 5’ tRNA halves compared to non-hematopoietic tissues and are suggested to function as immune signaling molecules [56, 57]. Consistent with these observations, we found that most transcripts ranged from 22-24 nt in the skeletal muscle and 33-36 nt in the liver. This represents the expected size of 5’ tRFs and 5’ halves respectively (**Figure 2B)**. Since tRFs of this size (22-24 nt) resemble miRNAs, this could indicate a higher likelihood of association with AGO proteins for gene silencing in the skeletal muscle [58]. Because the average size of a miRNA is ~22nt, we were curious if any of the predicted tRF sequences aligned to known miRNAs. We retrieved the mature sequences of all annotated bovine miRNAs from miRbase and aligned the tRF sequences using blast+ v2.10.1 [59, 60]. We found that none of the tRF sequences perfectly aligned to any of the bovine miRNAs.

**Figure 2.**
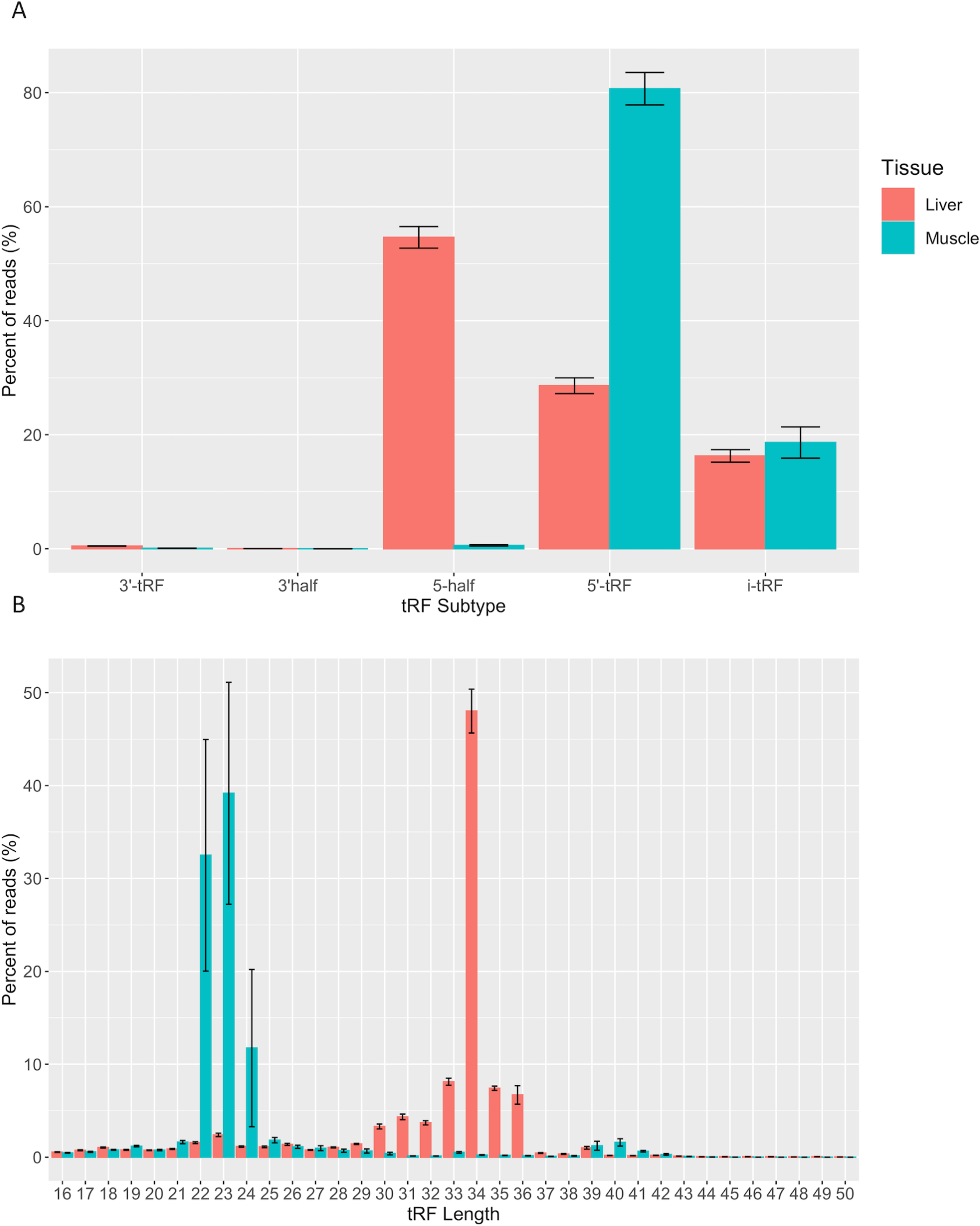
Quantitative analysis of tRF subtypes and size distribution. (A) Predicted tRFs were classified based on the region of the mature tRNA molecule that they are derived from across all samples (n = 10). (B) Predicted tRFs were also classified based on size across all samples (n = 10). All reads were categorized based on size or subtype and the y axis represents the percent of total CPM-normalized tRF transcript counts. The y axis sums to 100% for each tissue. Summary statistics were computed with the SummarySE function of the Rmisc package and standard error bars are shown in black.

We observed approximately 2.97% and 6.05% of the expressed transcripts in muscle and liver were derived from mitochondrial tRNAs. These observations suggest that MT-derived tRFs make a minor contribution to the tRFome. Consistent with a previous tRF study, the parental tRNA from which mitochondrial and nuclear tRFs originated, varied between muscle and liver [61]. For example, most MT-tRFs were derived from Lys^UUU^ in muscle and Asn^GUU^ in liver, while most nuclear tRFs were derived from initiator Met^CAU^ (iMet^CAU^) in muscle and Gly^GCC^ in liver **(Figure 3A&B)**. This finding demonstrates the unique expression profiles for nuclear- and MT-derived tRFs in the muscle and liver, which could underlie tissue-specific biological processes. In addition, we observed certain tRNAs did not produce tRFs in any of the treatment groups in the muscle or liver (Ala^GGC^, Arg^GCG^, Asp^AUC^, Cys^ACA^, Gly^ACC^, His^AUG^, Ser^ACU^, SeCe^UCA^, Thr^GGU^, Tyr^AUA^, and MT-Ile^GAU^). We previously found 10 of these tRNAs (excluding MT-Ile^GAU^) to be transcriptionally silent across all treatment groups in muscle and liver [21]. Despite the annotation of these silent tRNAs in the bovine assembly, a previous report has illustrated that 9 of these genes (excluding SeCe^UCA^ and MT-Ile^GAU^) are reportedly missing from eukaryotic, bacterial, and/or archaeal species [62]. This could suggest a selective pressure on anticodon bias across species. As far as we know, there are no reports of any tRNA isodecoders that do not participate in tRF biogenesis. This may indicate that some tRNA species are more resistant to processing events, and is perhaps linked to tRNA modifications offering protection from cleavage [63–65]. Finally, Phe^AAA^ is an example of a previously identified silent isodecoder with detected tRF expression, suggesting that some tRNAs are transcribed and cleaved to solely give rise to unique tRF species [21].

**Figure 3.**
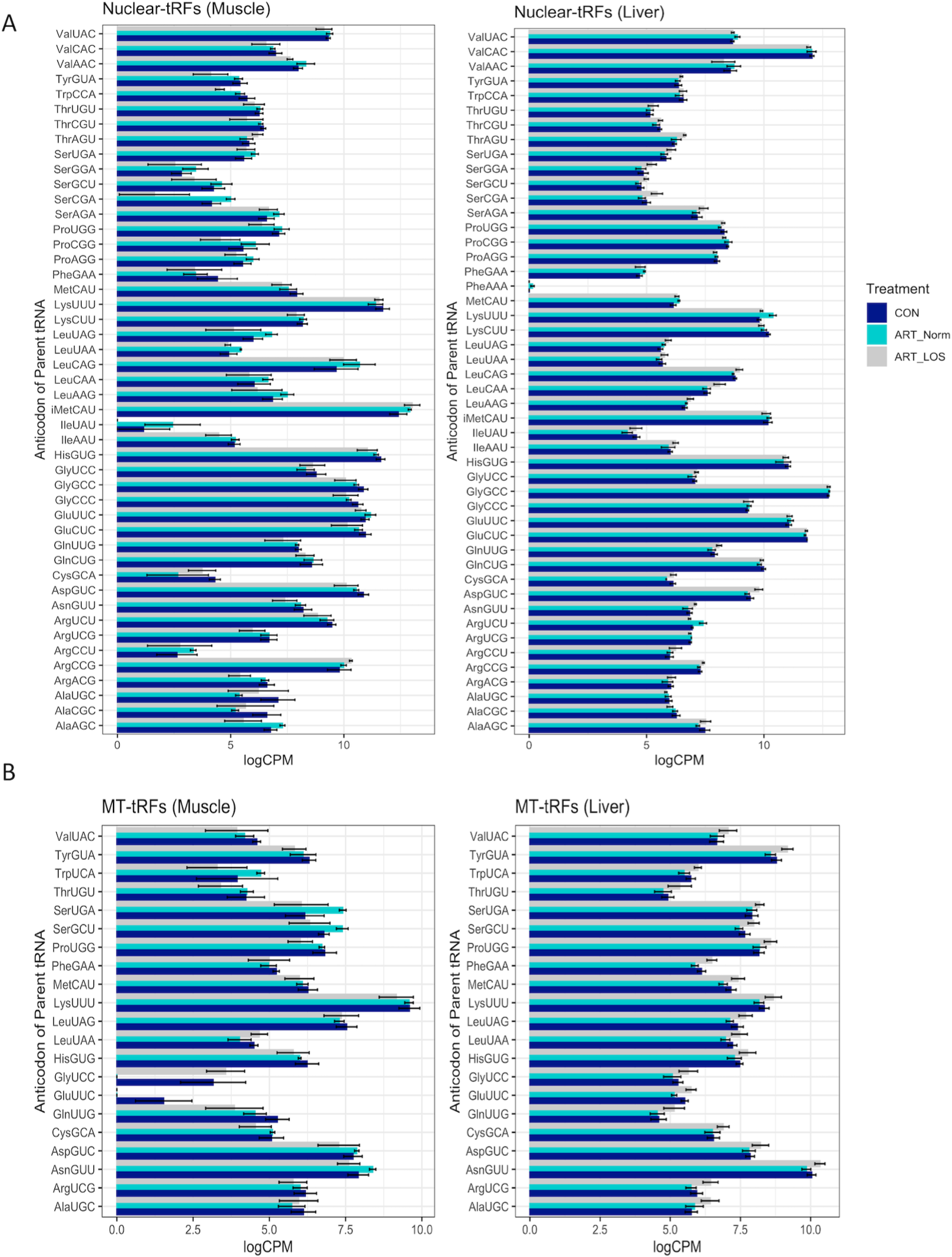
MT and nuclear tRF distribution. A bar graph depicting the levels of tRFs derived from (A) nuclear and (B) mitochondrial parental tRNAs across Control-AI, ART-Normal, and ART-LOS groups in muscle and liver tissue. Log transformed CPM values were used and each parental tRNA was grouped at the level of the anticodon. The SummarySE function was implemented to calculate the statistics of continuous variables by treatment group and standard error bars are shown for each anticodon in each treatment group.

### Data visualization by relative log expression and principal component analysis (PCA)

Relative log expression (RLE) plots were used to visualize the normalized tRF expression data across and within treatment groups **(Supplementary Figure 1)**. Most samples were constant although there was increased variation in ART-LOS #2 in the muscle **(Supplementary Figure 1A)** and ART-LOS #1 in the liver **(Supplementary Figure 1B)**. This is consistent with our previous work using tissue samples from the same ART-LOS individuals, in which genes in ART-LOS #1 in liver and ART-LOS #2 in muscle were expressed differently from other LOS individuals [19]. PCA plots show the clustering of individuals based on the normalized tRFs in muscle and liver **(Supplementary Figure 2)**. In the muscle, the Control-AI vs ART-Normal and ART-LOS vs ART-Normal cluster together, yet there is no clustering in the Control-AI vs ART-LOS groups **(Supplementary Figure 2A)**. Similarly, Control-AI vs ART-Normal and ART-Normal vs ART-LOS comparisons show clustering in the liver **(Supplementary Figure 2B)**. However, the PCA displaying all three treatment groups indicates that ART-LOS #2 clusters with the Control-AI group in the liver and away from other treatment groups in the muscle. Overall, we observed variation in tRF expression within treatment groups, particularly in ART-LOS individuals. This might be due to the nature of the syndrome, as certain LOS phenotypes differ in severity and are not always present [18]. Previous reports have suggested that tRFs may be less tightly regulated than other small RNAs, due to their larger abundance and the ability of each tRF to originate from several tRNA genes [66, 67]. However, several studies demonstrate that certain tRFs describe underlying mechanisms in cellular states and disease progression [30, 68]. Although overgrowth is one of the most common characteristics of LOS, liver tumor predisposition is variable and the classification of LOS fetuses based on body weight alone likely introduced a preference for tRF dysregulation in the muscle.

### Identification of differentially expressed (DE) tRFs

We used EdgeR v 3.24.3 to conduct differential expression analyses of the normalized tRF read counts in muscle and liver tissue [43]. We conducted DE analysis across three comparisons: Control-AI vs ART-LOS, Control-AI vs ART-Normal, and ART-Normal vs ART-LOS. For Control-AI vs ART-LOS, we identified 24 DE tRFs in muscle and detected no DE tRFs in liver **(Supplementary Table 2A)**. For ART-Normal vs ART-LOS, we identified 764 DE tRFs in muscle and 43 DE tRFs in liver **(Supplementary Table 2B)**. For Control-AI vs ART-Normal, we detected 196 DE tRFs in muscle and 44 DE tRFs in liver **(Supplementary Table 2C)**. We acknowledge that the low number of DE tRFs detected in the liver could be due to the classification of LOS fetuses based on body weight alone. We consistently saw higher numbers of DE tRFs in the muscle, which could suggest that the potential for gene targeting is higher in muscle tissue due to the recruitment of small RNAs that are similar in size to miRNAs **(Figure 2B)**. DE tRFs in Control-AI vs. ART-Normal and ART-Normal vs ART-LOS could suggest that tRF expression can be influenced by method of conception (AI vs ART) as well as syndrome development (ART-Normal vs ART-LOS). Additionally, heatmaps of DE tRFs in muscle and liver were produced to visualize the degree of up and down regulation across all individuals **(Supplementary Figure 3 & 4)**. The heatmap for muscle showed consistent DE expression across all treatment groups. There was much variation within Control and ART-LOS groups, which also could be explained by fewer DE tRFs detected in that comparison. The heatmap for liver displayed little consistency in expression between treatment groups, which could be due to the assignment of treatment group based on weight.

### Mature tRNAs are tightly regulated for non-canonical functions

Previously generated data from our study characterizing tRNA expression profiles in Control-AI, ART-Normal, and ART-LOS individuals was used in order to better understand the relationship between mature tRNA expression and tRF abundance [21]. Due to the high levels of sequence conservation across tRNA species, MINTmap can identify numerous parental tRNA sources for a single tRF. In an effort to not exclude any parent tRNA source, the counts for each tRF were divided by the number of tRNAs it was predicted to be derived from. In order to determine if there was an association between tRNA and tRF abundance, we performed a Pearson correlation analysis between tRNA and tRF expression. We found the Pearson correlation coefficients were 0.47 and 0.6 (p-value ≤ 0.05) for the muscle and liver respectively **(Figure 4A&B)**. One explanation for these moderately positive correlation coefficients is that selective transcription of tRNA genes can bias the availability of certain mature tRNAs and ultimately the population of tRFs. These findings agree with a previous study, which reported tissue-specific modulation of tRNA transcription to support its dual function in translation as well as gene regulation by tRFs [35]. This data demonstrates that tRF expression is non-random and dependent on the availability of highly regulated tRNA molecules. We acknowledge that the redundancy of tRNA genes and difficulties in efficient sequencing remains a major challenge in tRNA studies.

**Figure 4.**
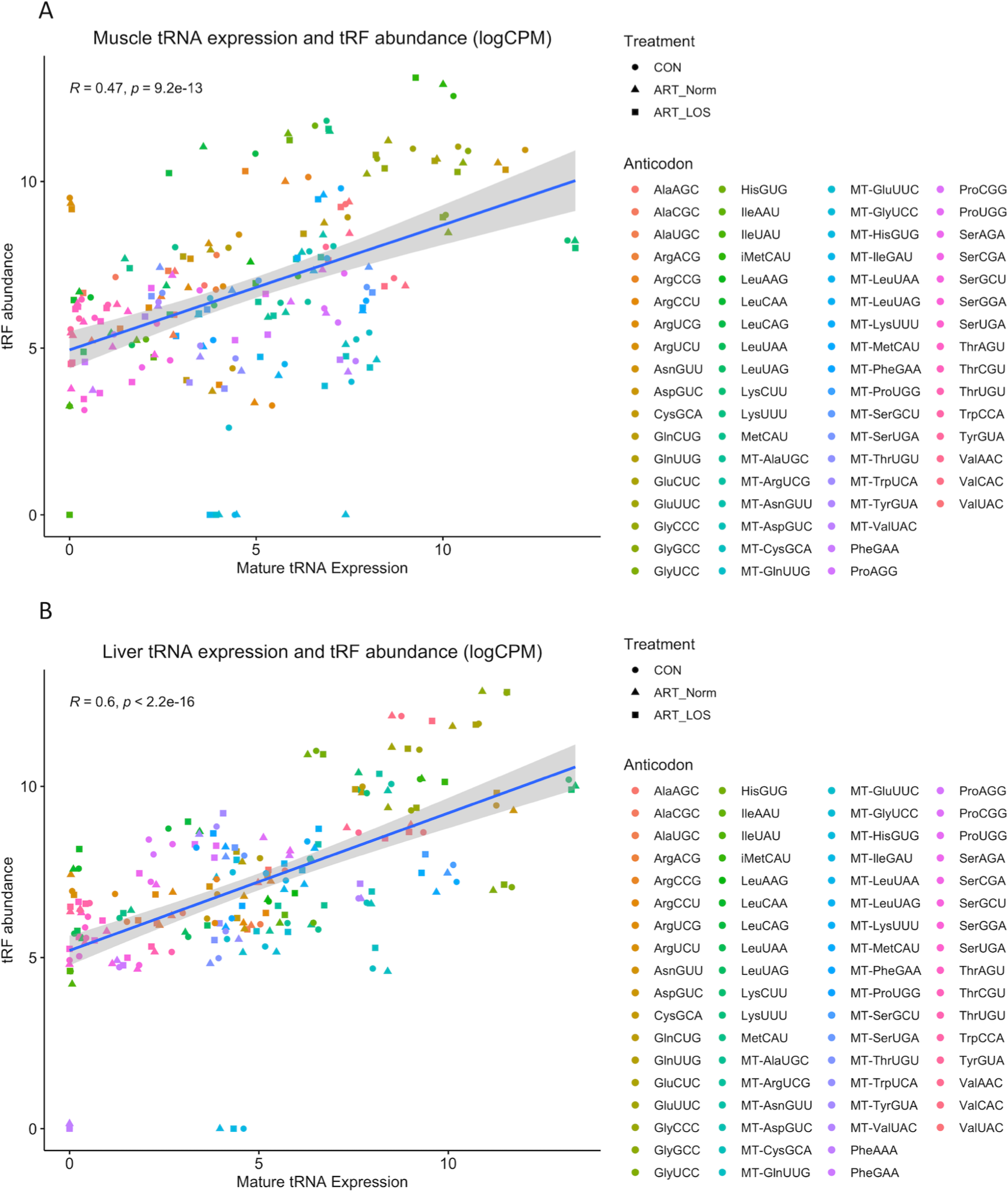
The relationship between muscle mature tRNA expression and tRF abundance. A scatterplot showing the correlation between tRNA (x-axis) and tRF expression (y-axis) in (A) muscle and (B) liver. Counts for the tRNA and tRF dataset were CPM normalized and log-transformed. tRNA species with no detected expression in any treatment group from either dataset were not included. The Pearson’s product moment correlation coefficient was used to determine the relationship between tRNA and tRF expression. Linear regression lines were added using the geom_smooth function of ggplot2 with the linear model argument.

### DE tRFs target transcripts in LOS individuals

Target prediction was done via miRanda with the sequences of the DE tRFs as well as the 3’ UTR sequences of all expressed protein-coding genes. The RNAseq datasets from a previous LOS study were used in order to identify DE mRNA transcripts and 3’ UTR sequences were retrieved from Ensembl Release 98 for the ARS-UCD1.2 reference genome [19, 45]. DE mRNA targets with an inverse relationship to DE tRFs were overlapped for each treatment group comparison and combined in order to generate candidate gene lists for each pairwise comparison. Pairwise comparisons were used for target prediction and enrichment. However, Control-AI vs ART-LOS in the liver was not used for further analysis because there were no statistically significant DE tRFs identified. R packages topGO v2.42.0 and KEGGREST 1.30.1 were used in order to identify functionally enriched biological processes, molecular functions, and pathways of all candidate target genes. In addition, g:Profiler was used to perform an analysis of human phenotype ontology (HPO) in order to identify enriched genes that are associated with phenotypic abnormalities in human disease [52]. There was no significant KEGG pathway or HPO enrichment in Control-AI vs ART-LOS in the muscle tissue. This is likely due to the low number of differentially expressed tRFs (24 DE tRFs predicted). Our enrichment analysis identified several affected biological processes, molecular functions, and signaling pathways between ART-Normal and ART-LOS groups in the muscle **(Figure 5A)** as well as abnormalities related to the targeted genes **(Figure 5B)**. Certain enriched HPO terms in the muscle were related to phenotypes often observed in LOS, such as hernia of the abdominal wall and abnormality of limb bone/skeletal morphology **(Figure 5B)**[69]. The full outputs for all performed enrichment analyses can be found in **Supplementary Table 3**.

**Figure 5.**
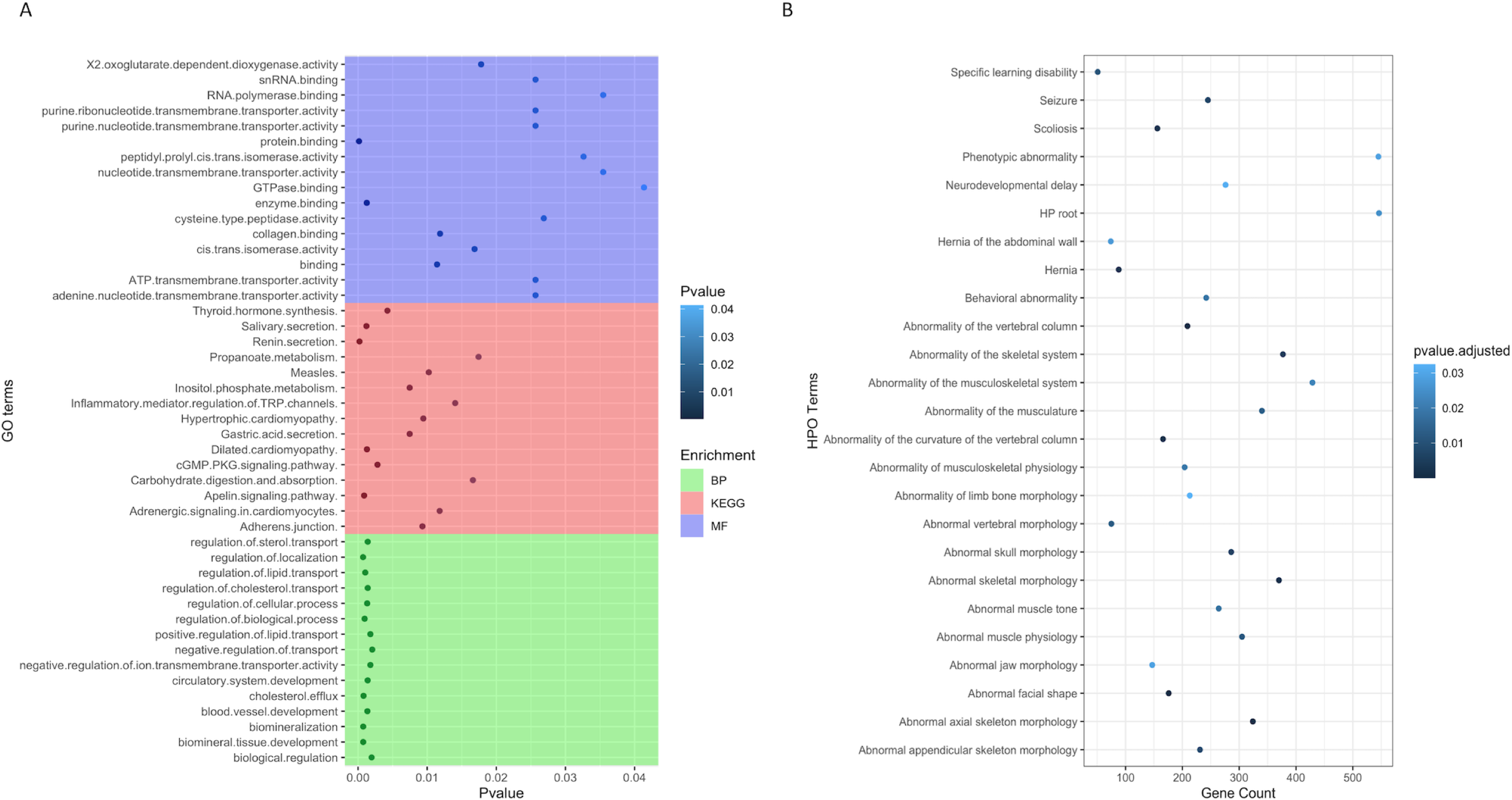
Biological processes, molecular functions and KEGG pathway enriched for target genes. (A) Subset of the affected biological processes, molecular functions, and KEGG pathways associated (p-value ≤ 0.05) with gene targets in ART-Normal vs ART-LOS muscle. Gene ontology (GO) enrichment tests were performed with TopGO using the classic algorithm with the fisher test. The pathway function of the KEGGREST package was used applying the Wilcoxon rank-sum test to calculate significantly enriched pathways. (B.) The top 25 enriched Human Phenotype Ontology (HPO) terms for gene targets in ART-Normal vs ART-LOS muscle. HPO enrichment was determined using g:Profiler with the g:SCS threshold significance criterion and HPO terms with an adjusted p-value ≤ 0.05 were considered significant. The number of genes enriched in an HPO term is shown on the x axis.

In the liver tissue, several GO terms associated with metabolic processes were enriched in Control-AI vs ART-Normal and ART-Normal vs ART-LOS (e.g. carbohydrate derivative metabolic process and glycoprotein metabolic process). In addition, there was an enrichment of genes related to immune response in both comparisons, such as regulation of phagocytosis, regulation of lymphocyte differentiation, and regulation of T cell differentiation **(Supplementary Table 3)**. Previous reports in other species suggest immune cells and inflammatory responses, such as phagocytosis, have a role in the progression of tumor development [70–72]. We also found that both the Wnt and cGMP-PKG signaling pathways were targeted in the liver of ART-LOS individuals. Of the enriched genes, *RACK1* and *MAPK3* were both upregulated in ART-LOS liver tissue and enriched in the Wnt signaling pathway and the cGMP-PKG signaling pathway respectively. *RACK1* is known to negatively regulate the Wnt signaling pathway, yet activate Sonic hedgehog (Shh) signaling [73, 74]. Activation of Shh signaling has been implicated as a potential prognosis predictor in human hepatocellular carcinoma and upregulation of *RACK1* in lung cancer correlates with metastasis and tumor differentiation [75, 76]. Our previous LOS study reported microRNAs targeting genes in the Wnt Signaling pathway as well, suggesting complementary mechanisms affecting gross regulators of LOS development [38]. The expression of *MAPK3* has been implicated in several cancer types, in which upregulation of *MAPK3* correlates with tumor recurrence and poor prognosis [77–79]. As previously mentioned, both ART-Normal and ART-LOS groups had enrichment of processes related to tumor formation. This could be due to the variability in the presence of LOS phenotypes and the assignment of individuals to a treatment group based on weight, suggesting both ART-Normal and ART-LOS could have increased chances of tumor development in liver.

In the muscle, gene targets were enriched in GO terms related to the regulation of biological process, cell cycle regulation, and tissue-specific developmental processes **(Figure 5A; Supplementary Table 3)**. We found *SMAD1* was enriched in the regulation of biological and cellular processes and was downregulated in ART-LOS individuals. *SMAD1* belongs to a family of anti-differentiation transcription factors that are critical to the bone morphogenetic protein pathway, which regulates muscle mass and regeneration [80]. The inhibition of *SMAD1* by microRNAs results in the promotion of skeletal muscle differentiation and regeneration [80, 81]. Furthermore, *BMI1* was upregulated in the muscle of ART-LOS individuals. Overexpression of *BMI1* in mouse mesenchymal stem cells causes an increase in body size, weight, length of tibiae, and width of the cartilaginous growth plate [82]. In addition, *RAI1* was downregulated in ART-LOS individuals. Changes in *RAI1* dosage can have significant impacts on growth and development. For example, overexpression of *RAI1* can result in extreme growth retardation, whereas haploinsufficiency of *RAI1* causes increased weight and fat deposition [83–85]. *RAI1* was enriched in the HPO term, abnormal appendicular skeleton morphology, suggesting that it could be responsible for phenotypic abnormalities in bovine (**Figure 5B**; **Supplementary Table 3**). According to our analysis and previous result, we confirmed the dysregulation of genes known to be associated with overgrowth: *IGF2R*, *GNAS*, *DNMT3A*, and *CDKN1C* [19]. *IGF2R* was downregulated in ART-LOS muscle and low levels of *IGF2R* has been identified in both bovine and ovine fetal overgrowth due to IVF [9, 86]. *GNAS* was upregulated in the ART-LOS muscle, which is consistent with hypomethylation of the *GNAS* loci that has been observed in Beckwith-Wiedemann syndrome [87]. Although mutations in *DNMT3A* are typically associated with overgrowth, we observed downregulation of this gene [88]. This could indicate that tRFs are capable of targeting DNA methyltransferases and modulating DNA methylation imprinting. In addition, a study using a mouse model found that embryos with *CDKN1C* deficiency can mimic phenotypes of BWS, such as overgrowth and abdominal wall defects [89]. Similar to this report, we observed downregulation of *CDKN1C* in ART-LOS individuals [89, 90]. Finally, we identified that *IGF1* was upregulated in ART-LOS. Shi and colleagues determined that the inhibition of a *FBXO40*, a negative regulator of *IGF1* signaling, resulted in elevated *IGF1* levels as well as increased body size and muscle mass in mice [91]. A list of tRFs that targeted the described genes above is shown in **Supplementary Table 3**. Although our analysis was limited to protein-coding genes for target prediction, we must recognize that long non-coding RNAs could also be regulated by small regulatory RNAs, such as miRNAs, and should be considered for future investigation [92].

Mammalian phenotype enrichment analysis (MamPhEA) based on mutant mouse phenotypes further revealed that tRF-regulated genes in muscle and liver tissue were associated with traits observed in LOS **(Figure 6)**. In muscle, genes were enriched for abnormal birth body size and abnormal skeleton morphology **(Figure 6A)**. In liver, genes were enriched for increased tumor growth/size and altered tumor pathology **(Figure 6B)**. Full results can be found in **Supplementary Table 3**.

**Figure 6.**
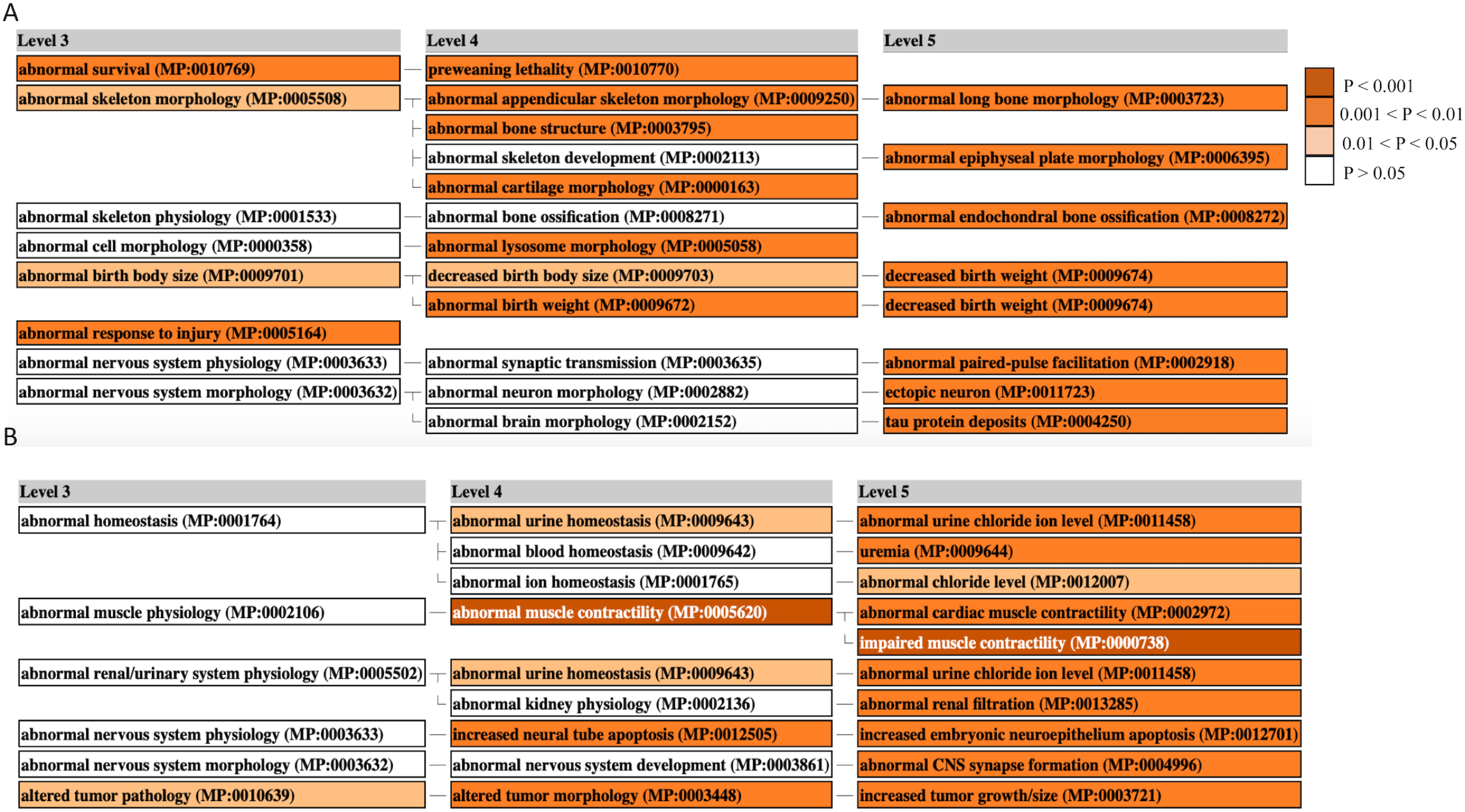
Mammalian Phenotype Enrichment Analysis. Enriched MGI phenotypes in ART-Normal vs ART-LOS A) Muscle and B) Liver. MamPhEA outputs shows enriched phenotypes that are hierarchically structured, representing mutant phenotypes enriched with more DEGs than by chance. Level 3 shows a general phenotype class and Levels 4/5 show detailed phenotype terms nested within the Level 3 class. Enrichment was determined using the Fisher’s exact test and phenotypes with a p-value ≤ 0.05 were considered significant. Enriched terms are shown in shades of orange.

## Conclusion

Overall, these data sets demonstrate that tRFs are commonly found in the muscle and liver tissue of Control-AI and ART-conceived individuals. Despite a moderate amount of variation in expression, we detected DE tRFs that may target pathways related to tumor progression or overgrowth. These outcomes provide deeper insight into the epitranscriptomic alterations that occur in ART-LOS individuals. This study is the first to examine the effect of altered tRNA availability on the differential expression of tRFs and its relationship to overgrowth syndrome.

## Abbreviations

ART: Assisted reproductive technologies
LOS: Large Offspring Syndrome
BWS: Beckwith-Wiedemann Syndrome
MOG: myelin oligodendrocyte
tRFs: tRNA-derived fragments
MT: mitochondrial
DE: Differentially expressed
DMRs: Differentially methylated regions
Pre-tRNAs: precursor tRNAs
i-tRFs: Internal-tRFs
RISC: RNA-induced silencing complex
AI: artificial insemination
RIN: RNA integrity number
PCR: Polymerase Chain Reaction
PCA: Principal component analyses
RLE: Relative log expression
FDR: False discovery rate
GSEA: Gene set enrichment analysis
HPO: Human Phenotype Ontology
MamPhEA: Mammalian phenotype enrichment analysis
SHH: Sonic hedgehog
IVF: In vitro fertilization
CDC: Center for Disease Control and Prevention
SRA: Sequence Read Archive
RNA POL III: RNA Polymerase III
TSEN Complex: tRNA Splicing Endonuclease Complex
AGO1-4: Argonaute 1-4
XPO5: Exportin-5
RanGTP: GTP-bound Ras-related nuclear protein
Nsun2: NOP2/Sun RNA methyltransferase family member 2
Dnmt2: DNA methyltransferase 2
NPC: Nuclear Pore Complex
ARS: aminoacyl-tRNA synthetase
ANG: Angiogenin
Mt-DNA: Mitochondrial DNA
RISC: RNA-induced silencing complex
eIF4A: eukaryotic translation initiation factor 4A
eIF4G: eukaryotic translation initiation factor 4G

## Data availability Statement

The Small RNA data for this project is deposited in FASTQ format to the NCBI Sequence Read Archive database (SRA) under the BioProject accession numbers PRJNA480853 and PRJNA876238. Additional RNAseq data is available at the NCBI Gene Expression Omnibus (GEO # GSE213525).

## Author Contributions

AKG carried out the experiment, sequencing library preparation, statistical data analysis, data interpretation, and the writing/revision of this manuscript. YL contributed to the data analysis and interpretation and revision of the manuscript. RMR provided tissue samples, assisted in conceptual design, contributed to data interpretation, and revision of the manuscript. DEH conceived the study, designed the experiment, was involved with the interpretation of the results and in the writing/revision of the manuscript. All authors read, revised, edited, and approved the final manuscript.

## Ethics Statement

All animal procedures were performed at TransOva Genetics by veterinarians, and all procedures were approved by their animal care and use committee (Protocol number -MRP2010-001) and were conducted in a manner conforming to Trans Ova Genetics policies and procedures and the Guide for the Care and Use of Laboratory Animals. The experiments were also conducted following the ARRIVE guidelines.

---

**Supplementary Figure 1:**
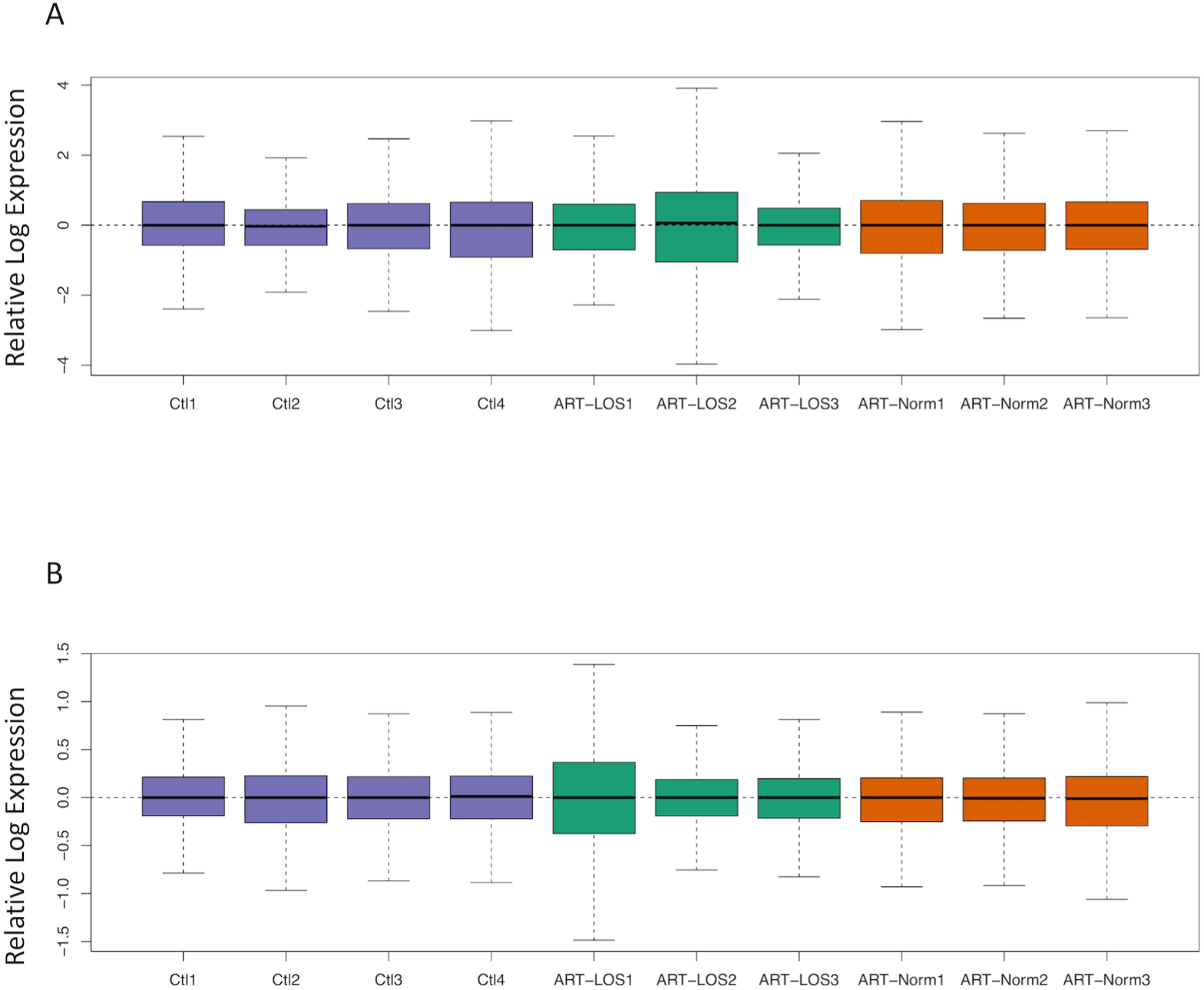
RLE plots to visualize variation of muscle tRFs across all treatment groups after normalization in (A.) muscle and (B.) liver.

**Supplementary Figure 2:**
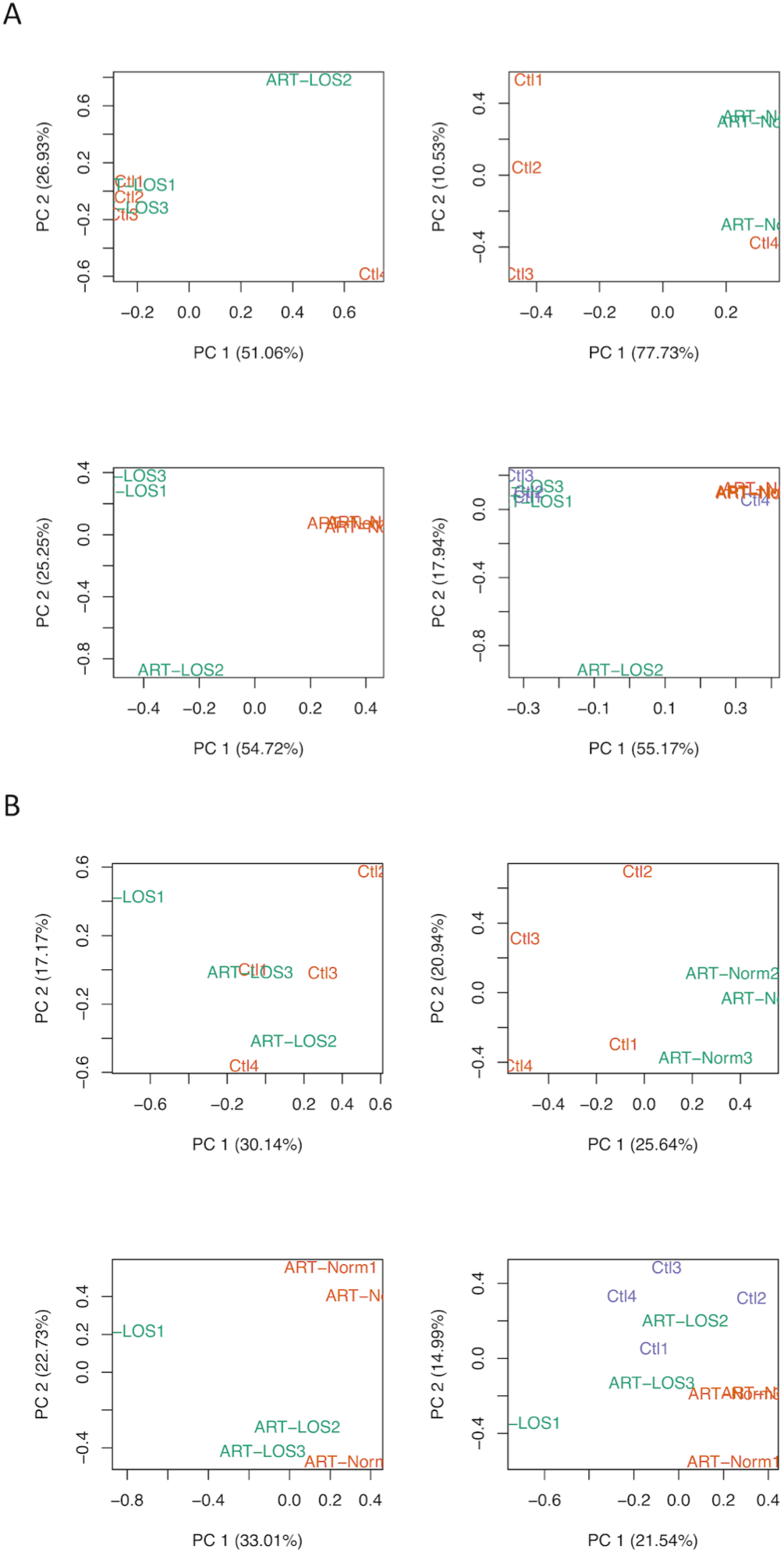
PCA plots of predicted tRFs in (A.) muscle and (B.) liver in different pairwise comparisons: Control vs ART-LOS, Control vs ART-Normal, ART-Normal vs ART-LOS, and all three treatment groups.

**Supplementary Figure 3:**
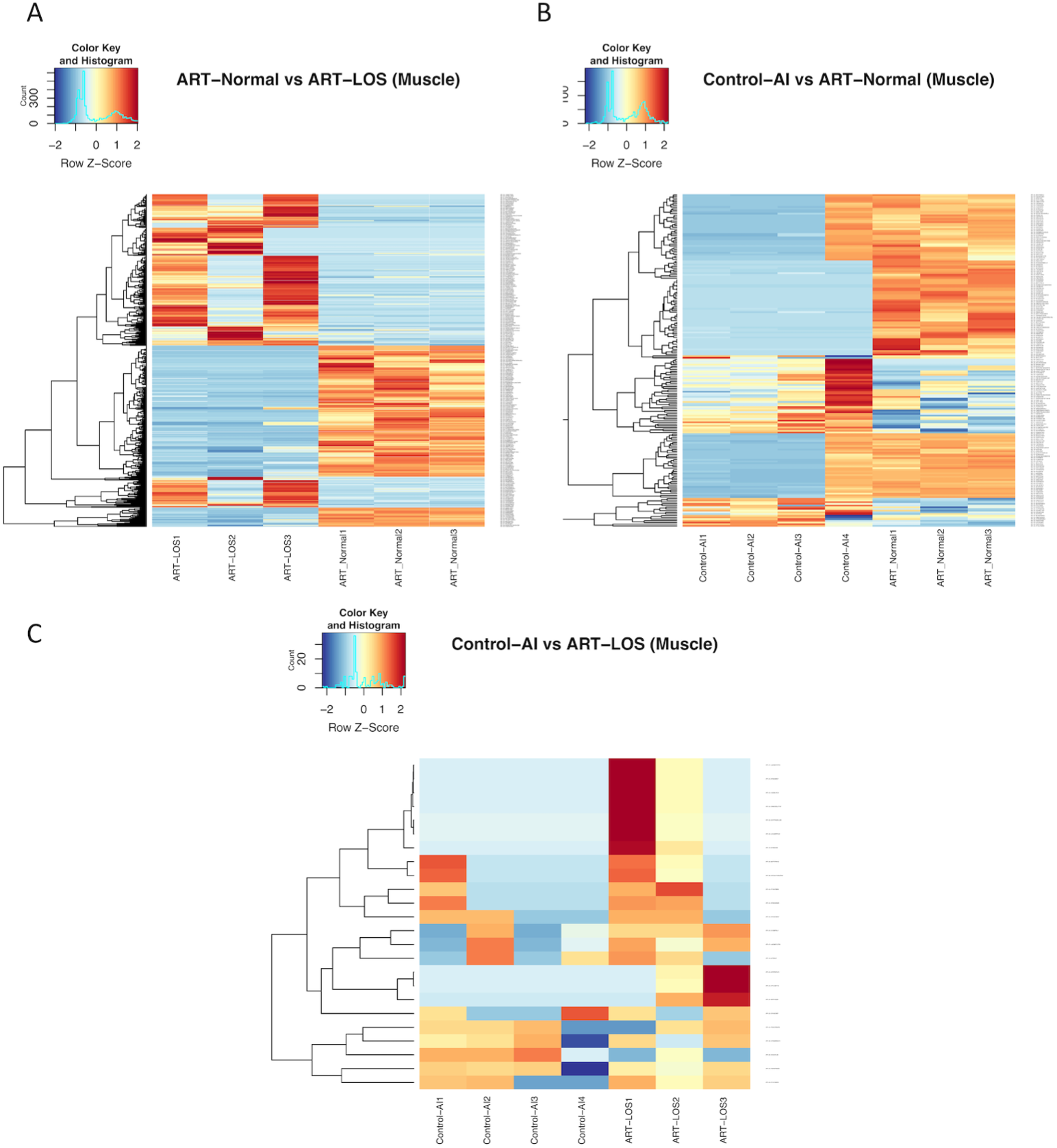
Heatmaps of differentially expressed tRFs in muscle. (A.) ART-Normal vs ART-LOS, (B.) Control-AI vs ART-Normal, and (C.) Control-AI vs ART-LOS.

**Supplementary Figure 4:**
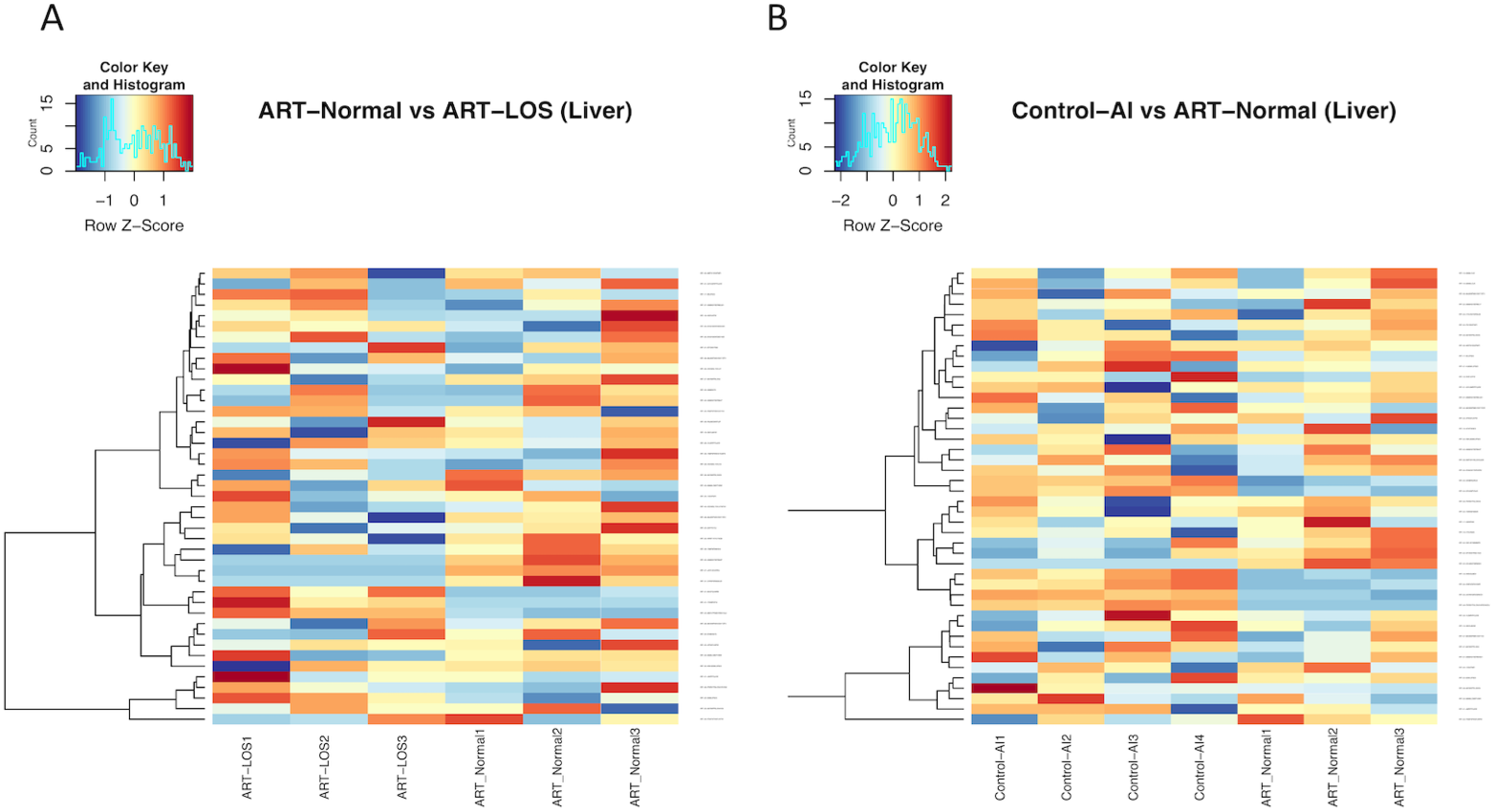
Heatmaps of differentially expressed tRFs in liver. (A.) ART-Normal vs ART-LOS and (B.) Control-AI vs ART-Normal.

